# A survey of neurophysiological differentiation across mouse visual brain areas and timescales

**DOI:** 10.1101/2022.01.21.476869

**Authors:** Saurabh R. Gandhi, William G. P. Mayner, William Marshall, Yazan N. Billeh, Corbett Bennett, Samuel D Gale, Chris Mochizuki, Joshua H. Siegle, Shawn Olsen, Giulio Tononi, Christof Koch, Anton Arkhipov

## Abstract

Neurophysiological differentiation (ND), a metric that quantifies the number of distinct activity states that the brain or its part visits over a period of time, has been used as a correlate of meaningfulness or subjective perception of visual stimuli. ND has largely been studied in non-invasive human whole-brain recordings where spatial resolution is limited. However, it is likely that perception is supported by discrete populations of spiking neurons rather than the whole brain. Therefore, in this study, we use Neuropixels recordings from the mouse brain to characterize the ND metric within neural populations recorded at single-cell resolution in localized regions. Using the spiking activity of thousands of simultaneously recorded neurons spanning 6 visual cortical areas as well as the visual thalamus, we show that the ND of stimulus-evoked activity of the entire visual cortex is higher for naturalistic stimuli relative to artificial ones. This finding holds in most individual areas throughout the visual hierarchy as well. For animals performing an image change detection task, ND of the entire visual cortex (though not individual areas) is higher for successful detection compared to failed trials, consistent with the assumed perception of the stimulus. Analysis of spiking activity allows us to characterize the ND metric across a wide range of timescales from 10s of milliseconds to a few seconds. This analysis reveals that although ND of activity of single neurons is often maximized at an optimal timescale around 100 ms, the optimum shifts to under 5 ms for ND of neuronal ensembles. Finally, we find that the ND of activations in convolutional neural networks (CNNs) trained on an image classification task shows distinct trends relative to the mouse visual system: ND is often higher for less naturalistic stimuli and varies by orders of magnitude across the hierarchy, compared to modest variation in the mouse brain. Together, these results suggest that ND computed on cellular-level neural recordings can be a useful tool highlighting cell populations that may be involved in subjective perception.

**Summary:** Advances in our understanding on neural coding has revealed that information about visual stimuli is represented across several brain regions. However, availability of information does not imply that it is necessarily utilized by the brain, much less that it is subjectively perceived. Since percepts originate in neural activity, distinct percepts must be associated with distinct ‘states’ of neural activity, at least within the brain region that supports the percepts. Thus, one approach developed in this direction is to quantify the number of distinct ‘states’ that the activity of the brain goes through, called neurophysiological differentiation (ND). ND of the entire brain has been shown to reflect subjective reports of visual stimulus meaningfulness. But what specific subpopulations within the brain could be supporting conscious perception, and what is the correct timescale on which states should be quantified? In this study, we analyze ND of spiking neural activity in the mouse visual cortex recorded using Neuropixels probes, allowing us to characterize the ND metric across a wide range of timescales all the way down from 5 ms to a few seconds. It also allows us to understand the ND of neural activity of different ensembles of neurons, from individual thalamic or cortical ensembles to those spanning across multiple visual areas in the mouse brain.

## Introduction

A key requirement for understanding the mechanistic origin of subjective, conscious visual perception is the ability to quantify visual experience based on neural activity. In the neuroscience of vision, for instance, regions of the primate brain that reflect information content of visual stimuli, such as edges, objects, faces etc. have been identified and extensively characterized (Rolls, 2000). But availability of information does not imply that it is necessarily utilized by the brain, much less that it is subjectively perceived (Brette, 2019; Buzsáki, 2019; Koch, 2004; Shimojo et al., 2001).

Consider, for instance, a conscious human observer watching a meaningful movie versus viewing an analog television displaying white noise or “snow.” (Fig. 1A). Both stimuli are constantly changing in time and have high diversity in space, so that, at the level of pixels on the screen, they both represent complex and highly dynamic patterns. However, for TV noise, the perceptual experience of an observer is low in complexity and remains approximately constant over time (TV noise is simply perceived as a more-or-less homogeneous, “noisy pattern” from moment to moment). In contrast, almost any scene from an engaging movie might change slowly, thus having lower temporal complexity, but is nonetheless more meaningful to the viewer, evoking distinct visual percepts over time. Since percepts originate in neural activity, each specific percept must correspond to a specific pattern of activity in the neural population that supports subjective visual perception, with a 1-to-1 mapping between the two (Boly et al., 2015).

**Figure 1:**
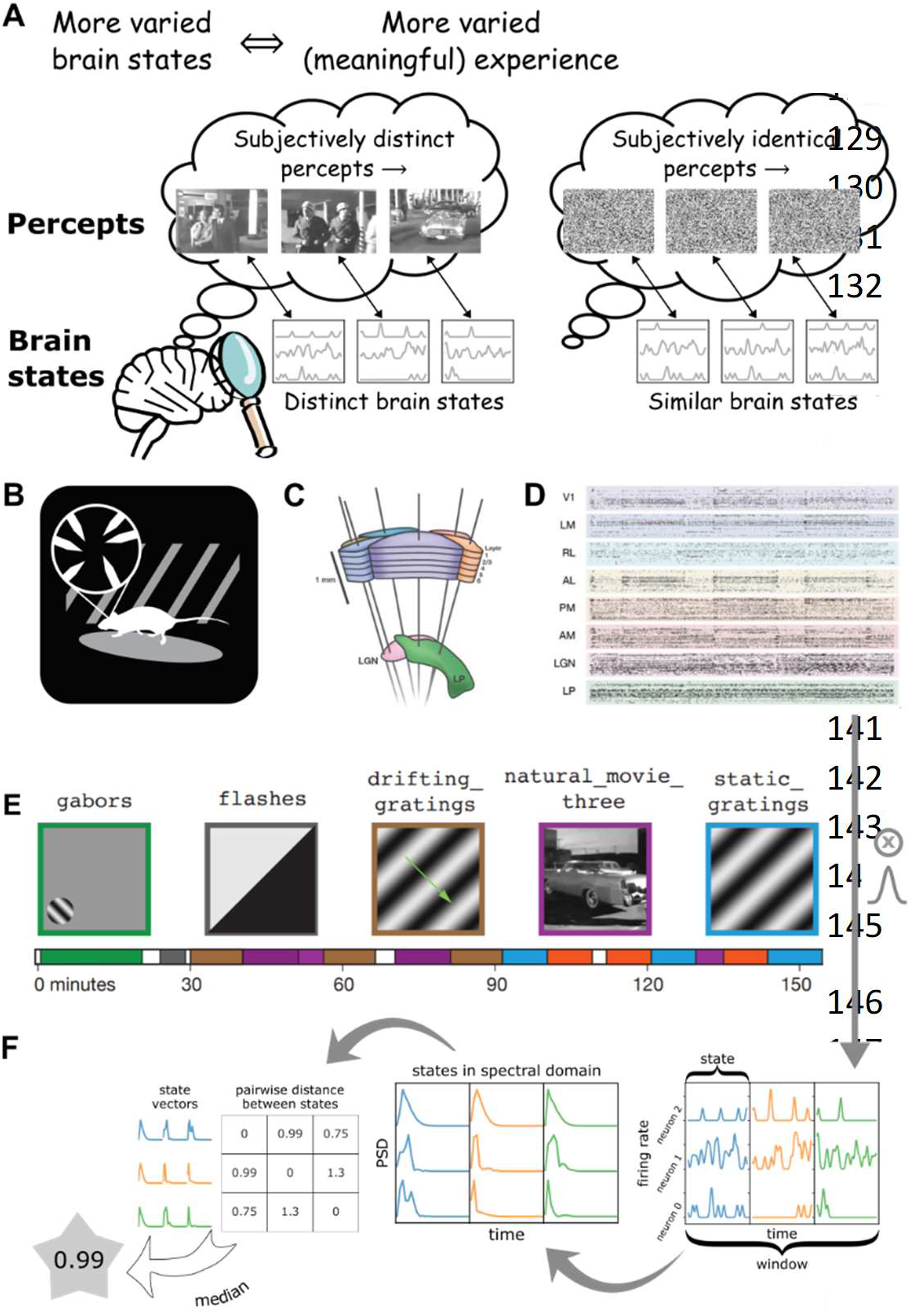
Quantifying ‘meaningfulness’ of subjective percepts using Neurophysiological Differentiation. **A.** Distinct subjective percepts must correspond to distinct physiological ‘states’ of the brain or the part of it that is the substrate for the experience. This can be captured by quantifying the size of state space explored by the appropriate brain region over a period of time. **B.** Schematic of the experiment. Mice a re head-fixed and free to run on a rotating platform. Visual stimuli are presented on a screen and responses are measured using 6 Neuropixels probes (**C**). **D.** Pre-sorted spiking activity simultaneously recorded from 6 visual cortical regions as well as thalamus and hippocampus were used for the analysis. **E.** Different classes of stimuli were presented including a control no-stimulus condition, simple artificial stimuli such as Gabor patches and full field flashes, more complex artificial stimuli such as static and drifting gratings and natural movies along with their time-shuffled versions. These stimuli span a wide range of stimulus complexity. **B-E** adopted with permission from (*Siegle et al., 2021*). **F.** The number of brain states are quantified using the Neurophysiological Differentiation (ND) metric. Spikes are convolved with a Gaussian; the resulting firing rate timeseries are divided into windows which are subdivided into states; states are quantified in terms of the PSD of each neuron’s activity with in each state concatenated into a single state vector; pairwise Euclidean distance is calculated between all states within a window; the median distance for a window is defined as the ND for the window.

In the example above, we would expect TV noise to result in relatively stable activity corresponding to the unchanging percept, whereas the movie scene would evoke temporally varying activity corresponding to each of the distinct percepts. Note that this will not be universally true for any neural population (for instance, it will not be true for retinal photoreceptors); here we refer only to a population that specifically supports subjective visual perception. Which neurons in the brain constitute such a population remains unknown at present. Thus, the richness of perceptual experience (‘meaningfulness of stimuli’) should correspond to the richness of neural activity (and not the richness of the stimulus), in the specific neuronal populations that are the physical substrate of the perceptual experience. This richness of neural activity has been quantified in several recent studies in terms of how many different ‘states’ the brain or a specific brain region goes through, called *neurophysiological differentiation* (Barttfeld et al., 2015; Boly et al., 2015; Gosseries et al., 2011; Hudetz et al., 2015; Mayner et al., 2022; Solovey et al., 2015; Synek, 1988).

A few different metrics of differentiation of neural activity, such as Lempel-Ziv complexity (Boly et al., 2015) or spectral differentiation (Mensen et al., 2018, 2017) have been proposed to infer the meaningfulness of visual stimuli to humans. Some of these metrics correlate with subjective reports from participants as to the ‘interestingness’, ‘understandability’ or ‘meaningfulness’ of the visual stimuli (Mensen et al., 2018). However, differentiation can be probed only at coarse spatial resolutions in human studies. To further our understanding of the relationship between differentiation of activity in specific brain regions and ‘meaningfulness’ of stimuli, we turn to studying differentiation at the cellular level in the mouse brain, leveraging readily accessible high-resolution and high-throughput recording techniques.

A recent calcium imaging study showed that the spectral differentiation of cellular fluorescent responses was higher for naturalistic stimuli (movies of predators and prey) compared to phase-scrambled versions of the same movies in two specific regions of the mouse brain, VISal and VISam, in layer 2/3. In layers 4 and 5 of the same areas and in all layers of the other areas studied (VISp, VISl, VISpm), the difference was non-significant (Mayner et al., 2022). In contrast, activity in any recorded visual cortical layer and region could be used to accurately decode stimulus type (naturalistic vs scrambled). While not a conclusive measure of perceived experience, the spectral differentiation metric was thus able to indicate with high specificity mouse brain regions that are potentially involved in perception in addition to the representation of stimulus information. Here, we apply a similar analysis to Neuropixels recordings from the mouse brain, allowing us to probe the metric across a wider range of spatiotemporal scales.

We first analyze the dependence of differentiation on the timescale of observation for single neurons as well as ensembles. We find that the differentiation of activity of single neurons does have an optimal timescale, close to the autocorrelation time of their firing rate; but this optimal timescale is shifted to under 5 ms for ensembles of neurons. We then fix the timescale and vary the composition of ensembles to understand how differentiation of response to different stimuli behaves in different regions of the mouse visual hierarchy. We find that in most (though not all) visual cortical areas, differentiation is higher for naturalistic stimuli compared to artificial ones. On the other hand, the ordering of response differentiation does not necessarily follow the ordering of the differentiation of the stimulus itself, suggesting that the metric we define is capturing more than just the information content of stimuli. We also show that depending on the behavioral state of the animal (whether it is running or stationary), differentiation of activity in individual cortical layers may or may not be modulated by the stimuli.

We repeat these analyses for mice performing a visual image change detection task to test the metric against a behavioral correlate of perception. Differentiation of activity of the entire visual cortex is higher for successful trials (hits) compared to failures (misses), though the difference was not significant in individual areas. These differences are also modulated by the behavioral state (running vs resting) of the animal.

Finally, we compare the differentiation metric between the mouse brain and artificial neural networks (ANNs) trained to perform a classification task. We see differences in magnitude as well as depth dependence of differentiation along the hierarchy in the two systems. Primarily, differentiation of activity in ANN layers does not correlate with how naturalistic the stimuli are.

Our results demonstrate that the differentiation metric applied to high resolution recordings can identify specific neural populations that appear to be sensitive to the expected perceptual differences in stimuli and correlates with a behavioral readout of perception. The results also highlight fundamental differences between biological brains and ANNs.

## Results

For the analysis presented here, we use the Visual Coding Neuropixels dataset (Siegle et al., 2021) published by the Allen Institute at www.brain-map.org; both Brain Observatory and Functional Connectivity stimulus sets were included in this analysis. For this dataset, mice were head-fixed but otherwise free to run on a rotating disk, while visual stimuli were displayed on a screen in front of them (Fig. 1B). Action potentials were recorded simultaneously from typically 6 Neuropixels electrodes consisting of 384 channels, sampled at 30 kHz, covering broad regions of the mouse cortical visual hierarchy (the primary visual cortex or area VISp, and the higher visual areas VISl, VISrl, VISal, VISpm, VISam; Fig. 1C) as well as thalamic regions (LGd, LP) and hippocampus. The dataset consists of pre-sorted spiking activity for 17,129 regular spiking (RS) and 3,910 fast spiking (FS) neurons (SNR > 2.5) across 58 recording sessions (distinct mice) and spanning the above-mentioned areas (~2,121 ± 663 RS and 470 ± 183 FS neurons per cortical area) (Fig. 1D).

Visual stimuli spanned a wide range of spatiotemporal complexity. Four broad classes of stimuli were presented:

(i) simple artificial stimuli such as full field flashes of white or black from grey and flashes of small Gabor patches; (ii) more complex artificial stimuli such as full field static or drifting gratings; (iii) naturalistic stimuli such as movie clips and (iv) temporally shuffled natural movies. A no-stimulus condition of a grey screen was also used (Fig. 1E). Each stimulus class was displayed in multiple blocks in ~3 hour recording sessions.

Here, we quantify the temporal complexity of spiking activity throughout the session using the spectral differentiation (differentiation for short) metric. Spike times are convolved with a Gaussian window (10 ms SD, cut off at ± 25 ms; see Methods) to obtain the firing rate of individual neurons as a function of time, sampled at 200 Hz. The firing rate (FR) timeseries is then divided into 3 s long non-overlapping windows *W*. Within each window, the FR is further typically split into 30 non-overlapping ‘state windows’ with a length of *S* = 100 ms. The corresponding state vector is obtained by computing the power spectrum of each neuron’s activity within that state window and then concatenating the spectra of all neurons into a single vector. Differentiation at a given time is defined as the median Euclidean distance between all pairs of state vectors within the window centered around that time (Fig. 1F, see Methods).

Since differentiation is defined using the Euclidean distance, the metric is not additive, i.e., differentiation of activity for a combination of ensembles is not the sum of differentiation for individual ensembles. This is because Euclidean distance between two vectors is not the sum of distances in orthogonal subspaces. Thus, while the differentiation metric primarily quantifies temporal complexity, it also accounts for the heterogeneity among neurons in a non-trivial manner. The metric does not, however, take into consideration spatial structure.

To enable comparison of differentiation across different ensembles of neurons, we apply additional normalizations (see Methods, Fig. S1). Firstly, if the firing rate of all neurons is scaled by a constant factor, the amplitude of the power spectrum, distances between pairs of spectra, and thus, differentiation for the ensemble, are all scaled by the square of that factor, even though the temporal complexity is unchanged. Therefore, to account for potential differences in overall firing rates across ensembles, the firing rates are divided by the average firing rate of all neurons within the ensemble across the entire recording time. Secondly, each neuron contributes a fixed number of dimensions to the state space, equal to the length of the power spectrum (see Methods). To enable comparison across differently sized ensembles of neurons, we further divide differentiation by the square root of the number of neurons in the ensemble, to obtain a per-neuron normalized metric. These two normalizations account for the uncontrolled heterogeneity (number and mean firing rate of neurons) in the ensembles analyzed.

The length of the state window (state length or *S;* see Methods) determines the timescale at which neural activity is observed. In the next section, we study how differentiation changes with the state length.

### Dependence of differentiation on the temporal scale of observation

Before analyzing the neurophysiological results, we build intuition for the dependence of differentiation on the timescale of observation, by computing differentiation for a signal using different values of state lengths *S*, ranging from 10 ms to 3 s, and lengths of windows, *W*, over which the number of states is quantified (3 s, 9 s and 30 s). For all instances discussed below, the sampling rate of the signal is fixed at 200 Hz.

First, as a toy example, we use an artificial signal that takes random integer values drawn uniformly between 0 and 255, with a characteristic timescale of 600 ms (the value of the signal changes every 600 ms). As the state lengths increase above the characteristic timescale, variations in the signal within the state are averaged out, and the median pairwise distance between state vectors should decrease. For state lengths shorter than the characteristic timescale, the median distance between pairs of state vectors should remain constant. This intuition is borne out when we compute differentiation (Fig. 2A): it indeed remains constant for state lengths shorter than the autocorrelation time, and decreases as a power law for longer state lengths. It is important to note that this trend can be different for a sparse signal (i.e., a signal which, for a given *W*, is mostly constant within the window): if the signal is sparse, then as *S* is shortened an increasing number of states have identical activity. Therefore, the median pairwise distance between state vectors continues decreasing to 0. Thus, for a sparse signal (such as spiking activity of neurons), differentiation is expected to be optimal near the characteristic timescale of the signal and decrease for longer and shorter state lengths (Fig. 2A).

**Figure 2:**
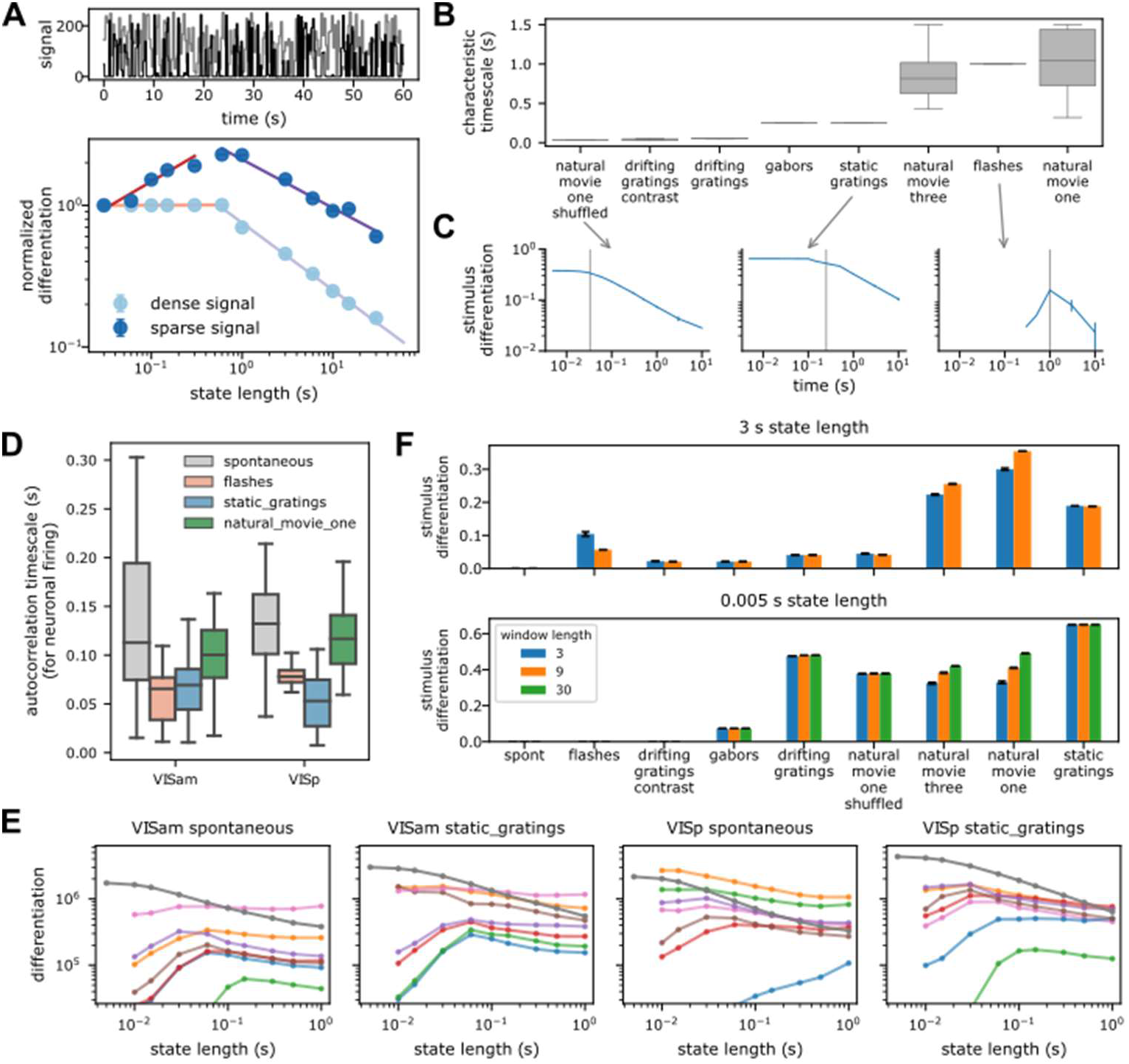
Dependence of differentiation on the temporal scale of observation. **A (top).** Toy example of a 1D timeseries with autocorrelation time of 600 ms (light: dense and dark: sparse signal). **A (bottom).** Differentiation of above signals as a function of state length S. Colored lines show power law fits. For S > autocorrelation time, differentiation decreases as a power law; for shorter S, differentiation remains constant for dense (light) but decreases for sparse signals (dark). **B.** Autocorrelation times (AC) of pixel values in the visual stimuli. Box plot shows distribution across all pixels. Vertical grey line indicates median of autocorrelation time from B. **C.** Stimulus differentiation (SD) vs S for three stimuli. Differentiation peaks (sparse signal) or plateaus (dense signal) around the AC of the signals (grey lines). Error bars are SD. **D.** Autocorrelation timescales of a subsample of neurons from a single session for responses to different stimuli **E.** Neurophysiological Differentiation (ND) of firing rate of individual neurons (colored lines) from two example areas and for two stimuli. Individual neurons have an optimal time at which ND peaks. Optimality shifts to timescale < 5 ms when ND is computed for all neurons from the respective areas (dark grey line). Error bars are SEM **F.** Differentiation for two extreme state lengths (top and bottom), for three example values of window lengths W in s (colors) for all stimuli. Differentiation is insensitive to W, but depends very sensitively on S, the timescale of observation (e.g., static gratings have the highest differentiation for S = 5 ms but movies have a higher differentiation for S = 3 s. Error bars are SD.

As a second example, we survey the differentiation of the visual stimulus itself. We treat each pixel of the stimulus as a ‘neuron’ and compute the ‘stimulus differentiation’ **(SD)** at different timescales, keeping *W* fixed at 3 s. Consistent with the non-sparse toy example above, for stimuli, SD remains constant up to the characteristic timescale (see Methods) and decreases with increasing state length as a power law beyond it (Fig. 2B, C). These stimuli consist of signals that vary substantially over the entire length of the window, and thus, differentiation plateaus at state lengths shorter than the characteristic timescale. Two stimuli – flashes and short drifting gratings – are sparse stimuli with activity concentrated in short intervals of time and the signal remains constant otherwise. For these two stimuli, differentiation has an optimum near the characteristic timescale and decreases for both shorter and longer state lengths, as expected.

Note that for a similar kind of activity (such as similar stimuli, or neural responses to similar stimuli), we expect the activity to occupy nearby regions in state space. Consequently, as the window size increases, the median distance between states should asymptotically approach a constant value, as we can observe in both the stimulus differentiation and response differentiation for an example experiment (Fig. S2). Therefore, for all of our analyses, we fix *W* at 3 s, a trade-off between longer windows where differentiation is close to the asymptotic value and shorter windows that facilitate computational tractability.

Next, we ask how the differentiation of responses of individual neurons (called neurophysiological differentiation, or **ND**) depends on the time scale. Consistent with the observations above, ND of activity of a large fraction of single neurons is maximized at some optimal timescale, corresponding to the characteristic timescale of their activity (Fig. 2D, E; see Methods). Roughly 27% (373 / 1,429 per cortical area) of regular spiking (RS, putatively excitatory pyramidal) cortical neurons show such an optimal timescale (~56% thalamic RS neurons), averaged across stimuli (Fig. S3). A slightly larger fraction (~42%; 113 / 274 per area) of fast spiking (FS, putatively inhibitory interneurons) cortical neurons have an optimal timescale. A majority of the remaining neurons have an optimal timescale longer than 1 s, which was not probed, while a few have very low firing rates, and thus zero ND at all state lengths.

The intrinsic timescale of correlations of neural activity increases along the anatomical visual cortical hierarchy in this same dataset (Piasini et al., 2021). The optimal timescale of ND for RS neurons is also shortest in the thalamic areas, and is longer, but surprisingly remains relatively constant (p > 0.05 for all stimuli) across the visual cortical hierarchy. The optimal timescale is shortest in layer 4 for RS neurons (Fig. S3). These modulations with laminar depth or cortical area are largely restricted to RS neurons, and are not as strong in FS neurons (p > 0.05 with respect to hierarchy except for Gabor stimuli). Optimal timescales for FS neurons are around 40-60 ms shorter than for RS neurons. Overall, optimal timescales for individual neurons vary within a factor of 2 across stimuli, ranging from 100 ms to 225 ms in cortical areas. They are not related to the optimal timescales for the corresponding stimulus differentiation either.

Finally, we ask how ND for an ensemble of neurons depends on the timescale. Interestingly, although single neurons have an optimal ND timescale, this optimality is essentially lost for ensembles (Fig. 2E, S5): ND increases at shorter and shorter timescales down to 5 ms (with a sampling rate of 200 Hz, we cannot consider state lengths shorter than 5 ms; since we do not find an optimal timescale within this range, we refer to this as ‘loss of optimality’ subsequently). Although individual neurons within the ensemble might have a similar characteristic timescale, the sparsity of firing and jitter or offsets in spike times are the likely causes for increasing ND at very short timescales. We verify this hypothesis by computing ND for an ensemble of virtual neurons, constructed by taking the spiking activity of a single neuron and adding random jitter or offset to the spike times. For neurons with a high firing rate, very small ensembles of 3-4 virtual neurons already show loss of optimality. For low firing rate neurons, the optimality of ND moves to shorter timescales as the number of virtual neurons increases (Fig. S4) but remains above 10 ms. Thus, with an increasing number of neurons in an ensemble, sparsity of their collective activity is reduced, resulting in the decrease of the optimal times for ND of the ensemble activity.

Analysis of our empirical data shows that even for a small number of neurons, the ND optima shift into the sub-5 ms range. This underscores the important point that ND should not necessarily be expected to be maximal at a timescale relevant to the stimulus or the experience, unlike a quantity that would fully characterize conscious experience, which involves both differentiation of the experience and integration of its components (Tononi et al., 2016). Rather, ND should be considered a relative measure defined at a timescale chosen by the observer, which can be compared across conditions (stimuli, brain states, etc.) or, after using an appropriate normalization, across neural ensembles. The dependence of ND on the timescale of observation is affected by the characteristic timescale of the underlying signal, which can have significant consequences on the interpretation of differentiation results. For example, at short timescales, the stimulus differentiation of static gratings is higher compared to natural movies, but at longer timescales, the relationship inverts (Fig. 2F).

In the following sections, we study the dependence of ND on cortical areas, layers, and the complexity or relevance of the stimuli. Since single neurons, on average, show an optimal timescale of 100 ms, we fix this as the state length for subsequent analysis. This timescale is also appropriate given that it is the timescale at which the dynamics of subjective experiences are thought to occur (Del Cul et al., 2007; Herzog et al., 2020, 2016).

### Modulation of ND by different visual stimuli across brain areas

The Visual Coding Neuropixels dataset includes a wide variety of images and movies, with varying levels of stimulus differentiation (SD), with grey screen having zero differentiation, static gratings having the highest and natural movies being intermediate (for the 100 ms state length; Fig. 3A). In addition to SD, another factor that could influence the ND of responses is the relevance or meaningfulness of the stimulus to the animal. In a brain region whose activity determines perceptual experience, ND is expected to have a stronger correlation with meaningfulness of the stimulus than with SD. Although we cannot ascertain the actual meaningfulness of the stimuli in this dataset to the mouse, we can ascribe a putative meaningfulness or relevance in terms of how close the stimuli are to a naturalistic setting. For example, an unchanging grey screen is putatively the least relevant, artificial stimuli, unlikely to occur in a natural setting, such as gratings, a little more relevant, and naturalistic movie most relevant given the higher order spatio-temporal structure in movies that is absent in gratings.

**Figure 3:**
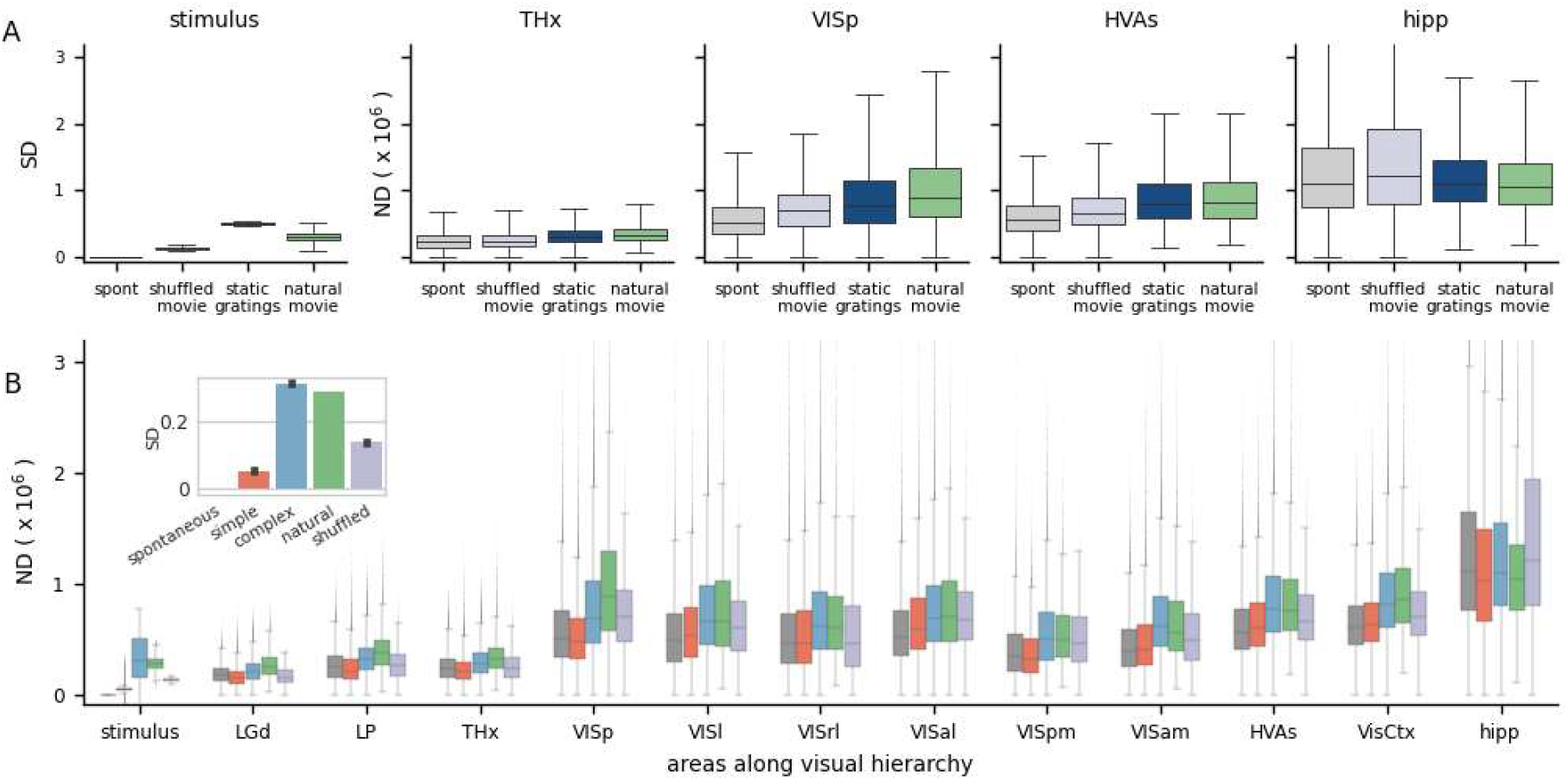
Modulation of differentiation by visual stimuli across brain areas. **A.** Stimulus differentiation (SD) for the 4 stimuli, computed for S = 100 ms, for stimulus pixels and neurophysiological differentiation (ND) (× 10^6^) for 4 neuronal ensembles; these do not simply reflect stimulus differentiation but putative relevance of stimulus – static grating stimuli (blue) are more differentiated than natural movie stimuli (green), but responses to movies are more differentiated than gratings in most brain areas (all pairwise differences are statistically significant). THx – all visual thalamic areas, HVAs – all higher visual areas, hipp – hippocampal areas **B.** ND increases going from thalamus to VISp but remains constant over the visual cortical hierarchy. Simple artificial stimuli (full screen flashes and Gabor patch flashes) evoke least ND, followed by complex artificial stimuli (gratings) and natural movies (green) typically evoke the highest ND in most areas. Interestingly, (a) time-shuffled movies sometimes evoke a higher or comparable ND as natural movies and (b) spontaneous activity sometimes has a higher ND than simple artificial stimuli. Hippocampal ND is not modulated by the stimulus type, consistent with it not playing a direct role in vision.

We compare the ND of neural responses to these stimuli in 4 regions: visual thalamus (THx), primary visual cortex (VISp), higher visual cortices taken together as an ensemble (HVAs) and, as a control, the hippocampus (hipp), which is not a visual region (Fig. 3A; due to the large number of neurons recorded, ND is statistically significantly different between all regions p < 0.001). Consistent with expectation, the response to grey screen (spontaneous activity) is least differentiated in all visual brain areas. This is not surprising given that it also has zero SD. Interestingly, however, even though static gratings are much more differentiated than natural movies, responses to gratings are less differentiated than those to movies in all three visual brain areas shown. Thus, the ND metric does not simply reflect the differentiation of the stimulus but correlates with the putative relevance of the stimulus to the mouse.

Next, we group the Visual Coding dataset into 5 broad categories with increasing putative meaningfulness - no stimulus (grey screen or spontaneous activity), simple artificial stimuli (full screen flashes and flashes of small Gabor patches), time-shuffled movies, complex artificial stimuli (static and drifting gratings) and natural stimuli (movies with higher order spatiotemporal correlations). SD for these stimuli is ordered differently than their putative relevance: spontaneous, simple artificial, shuffled, natural and finally complex artificial (Fig. 3B, inset). This sets up an interesting question of whether ND follows either of the two orderings (putative relevance or SD) or is entirely different. ND would follow putative relevance for ensembles encoding stimulus meaningfulness and SD for ensembles encoding information only. For areas not involved in vision, such as the hippocampus, we do not expect any particular ordering of ND.

We compute ND for ensembles of neurons restricted to individual areas along the visual hierarchy, or their combinations, such as neurons from all higher visual areas (HVAs), or the entire visual cortex (VisCtx) etc. (as noted earlier, ND is not an additive metric; ND of a combination of areas is not the same as the sum of ND from individual areas). Broadly, we find that ND increases going from thalamic areas to the cortex, but mean ND remains roughly constant over the hierarchy of visual cortical areas (Fig. 3B; p > 0.05 for all stimuli).

Across much of the visual hierarchy, putatively less meaningful stimuli (grey screen and simple artificial stimuli) evoke the least differentiated responses, complex artificial stimuli evoke more, and natural stimuli evoke the most differentiated responses (exceptions are VISrl and VISam, in which ND follows the same ordering as SD). The latter point is particularly interesting, since SD is higher for complex artificial than natural stimuli. The ND metric in most individual visual areas may therefore reflect the putative relevance of the stimuli rather than the pixelwise stimulus differentiation.

Note that the no-stimulus condition has a more differentiated response than simple artificial stimuli in a few areas (VISp, VISrl, VISpm). This observation is also consistent with the possibility that ND reflects a measure of perceptual experience rather than just the stimulus properties (see Discussion).

Finally, the hippocampus exhibits substantially higher ND than the other areas analyzed. This suggests a greater diversity of activity states in the hippocampus than in the visual cortex and thalamus, possibly reflecting a more narrow, specialized role of the latter structures dedicated primarily to visual processing. This does not mean, however, that the hippocampus is necessarily more involved in perception - only that the activity there is more temporally complex (i.e., occupies a larger volume of the state space). For identifying areas involved in conscious perception, it is not the magnitude of responses, but modulation of responses based on stimulus type, that is relevant. Importantly, hippocampal ND is uncorrelated with putative meaningfulness as well as stimulus differentiation, as expected given that broadly the hippocampus does not directly respond to visual stimuli.

While the results presented in this section are obtained for regular spiking neurons only, they hold for fast spiking neurons as well (Fig. S6). Although ND of fast and regular spiking neurons shows similar patterns, the overall magnitude of ND of cortical FS neurons is about half that of RS neurons, while thalamic FS neurons have similar ND as RS. Given the similarities, we restrict our analysis to RS neurons in subsequent sections.

### Modulation of ND by visual stimuli across cortical layers

To visualize the modulation of ND by visual stimuli for all areas and layers, we plot the difference of ND for all pairs of stimuli as a matrix (Fig. 4). The stimuli are ordered along the axes according to their putative relevance, so that positive differences (in red) indicate a positive relationship between the putative relevance and evoked ND (stars indicate significance: * p < 0.01; ** p < 0.001; *** p < 0.0001). Pairwise differences and significance are obtained by fitting a linear mixed effects model with fixed effects of layer, area and stimulus category, and random effect of mouse. Consistent with the previous section, this visualization reveals ND of entire areas is positively correlated with putative relevance for all areas except VISrl and VISam (Fig. 4, bottom row, bottomright cell: natural - complex artificial). Also as previously noted for entire areas (i.e., combining all layers), spontaneous activity is more differentiated than the responses to simple artificial stimuli in VISp, VISrl and VISpm (Fig. 4, bottom row, top-left cell).

**Figure 4:**
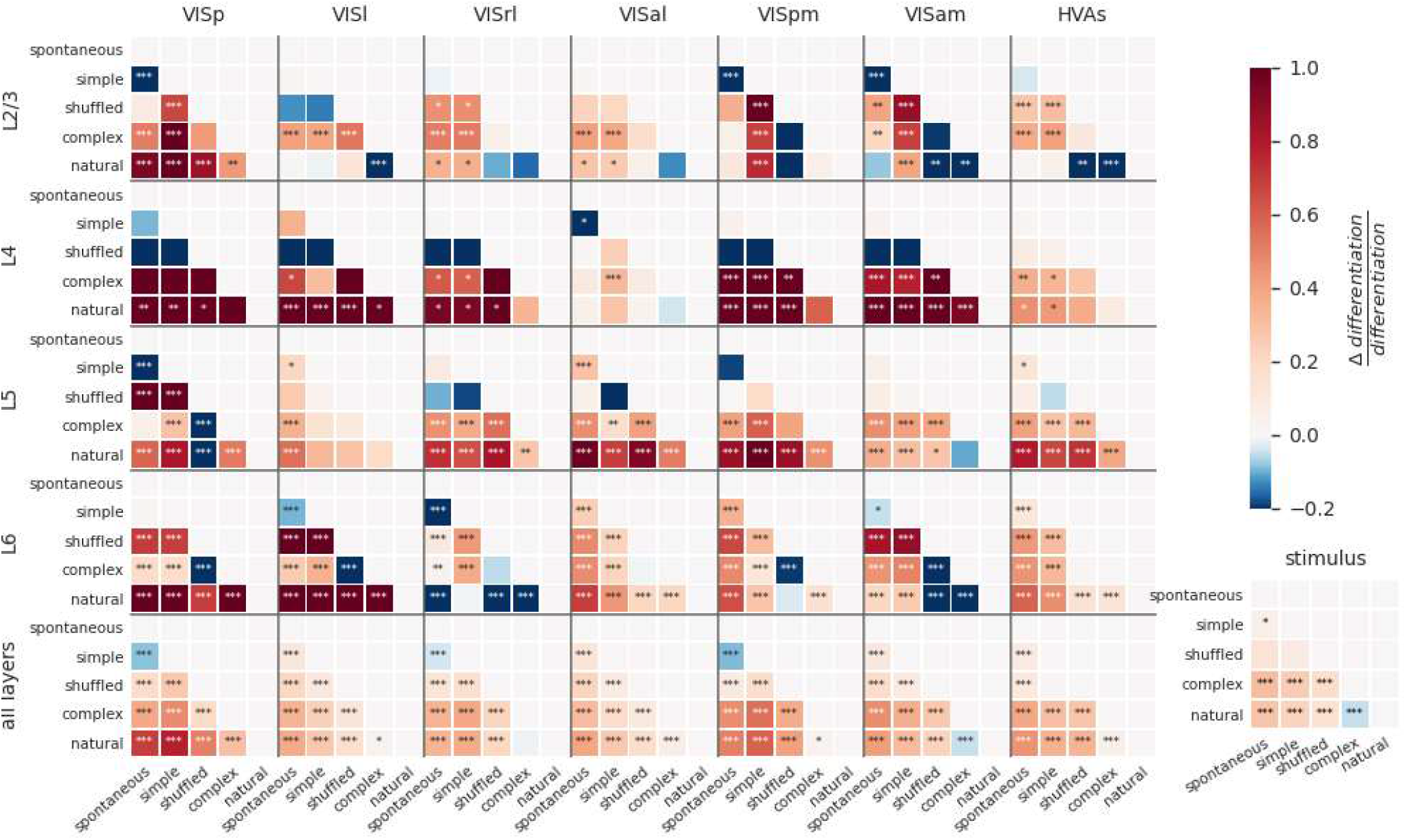
Modulation of differentiation by visual stimuli across cortical layers. Difference in ND of cellular responses to stimulus pairs for different areas and layers. Within each panel, each square shows difference of ND for y-axis and x-axis stimuli (red indicates y-x is positive; blue indicates y-x is negative). Stars indicate statistically significant differences (* p < 0.01; ** p < 0.001; *** p < 0.0001). L6 shows the most significant modulation of ND by stimulus, although most other areas also show significant modulation for several stimulus pairs. ND is consistent with putative meaningfulness in layers 4 and 5; but in layers 2/3 and 6, natural movies often evoke lower ND and shuffled movies evoke higher ND than other stimuli. This difference disappears when the analysis is restricted to when the mouse is running (Fig. S7).

We next investigate the layer-specific contributions to the ND of responses to different stimuli. Overall, within layers 4 and 5 of individual areas, differences in ND for most pairs of stimuli are broadly consistent with putative meaningfulness. Interestingly, ND of responses of layers 2/3 and 6 of higher visual areas does not follow this pattern, and natural movies tend to evoke lower ND compared to several other stimuli; shuffled movies tend to evoke higher ND than natural or complex artificial stimuli (see Discussion).

Depending on the behavioral state of the animal, the differences between layers become starker (Fig. S7). When the animal is running, the discrepancies between ND and putative meaningfulness in layer 6 are highly reduced compared to the resting state. Secondly, the overall statistical significance of modulation of ND by stimuli is much reduced in all layers except layer 6 in the running state.

Note that layers 2/3, 4, 5 and 6 have a comparable number of recorded neurons (52 ± 25, 44 ± 17, 73 ± 26 and 52 ± 24 respectively on average per mouse) while layer 1 has fewer (15 ± 9) and a more inconsistent number of neurons across experiments. Layer 1 is thus excluded from this analysis.

ND patterns in individual areas can be very different from those seen when the areas are combined. For example, shuffled movies evoke higher ND than complex artificial stimuli in layer 6 of all individual cortical visual areas (VISp, VISl, VISpm, VISam are significant). Yet, when all neurons are combined (L6 AllVis / HVAs), their ND is approximately equal for responses to complex artificial and shuffled stimuli (no significant difference), underscoring the non-additive nature of the ND metric.

Together, this shows that the differences between stimuli are reflected in differentiation of neural activity in all layers individually. However, the evoked ND is not necessarily consistent with putative relevance of the stimuli, as seen in layers 2/3 and 6. When all layers are taken together, however, the evoked ND is consistent with putative relevance, suggesting that it’s the larger ensembles of neurons across all layers that may be supporting or reflecting stimulus meaningfulness.

### ND during behavior

Although we show a correlation between the putative relevance of stimuli and ND of neural response, we cannot directly access the actual experience of the mouse - i.e., whether it even perceives the stimulus or not. Towards addressing this concern, we compute ND of responses to visual stimuli as the mouse performs a visual behavior task (Siegle et al., 2021).

In this task, water-deprived mice (n = 24) were head-fixed but otherwise free to run on a disk. A series of images was flashed in front of the mice, and the mice were trained to detect a change in the identity of the image by licking at a spout. Successful detection of change, called a hit, was rewarded with water. The mice did not get the water reward if they did not lick in response to the image change, called a miss (Fig. 5A). Neural activity was recorded using Neuropixels probes, yielding a total of 7638 neurons (318 ± 150 per mouse) with SNR > 2.5. For a mouse detecting an image change and responding, hits are likely correlated with successful perception and misses with lack of perception. We thus expect ND of spiking activity to be higher for hits compared to misses in areas representing experienced percepts.

**Figure 5:**
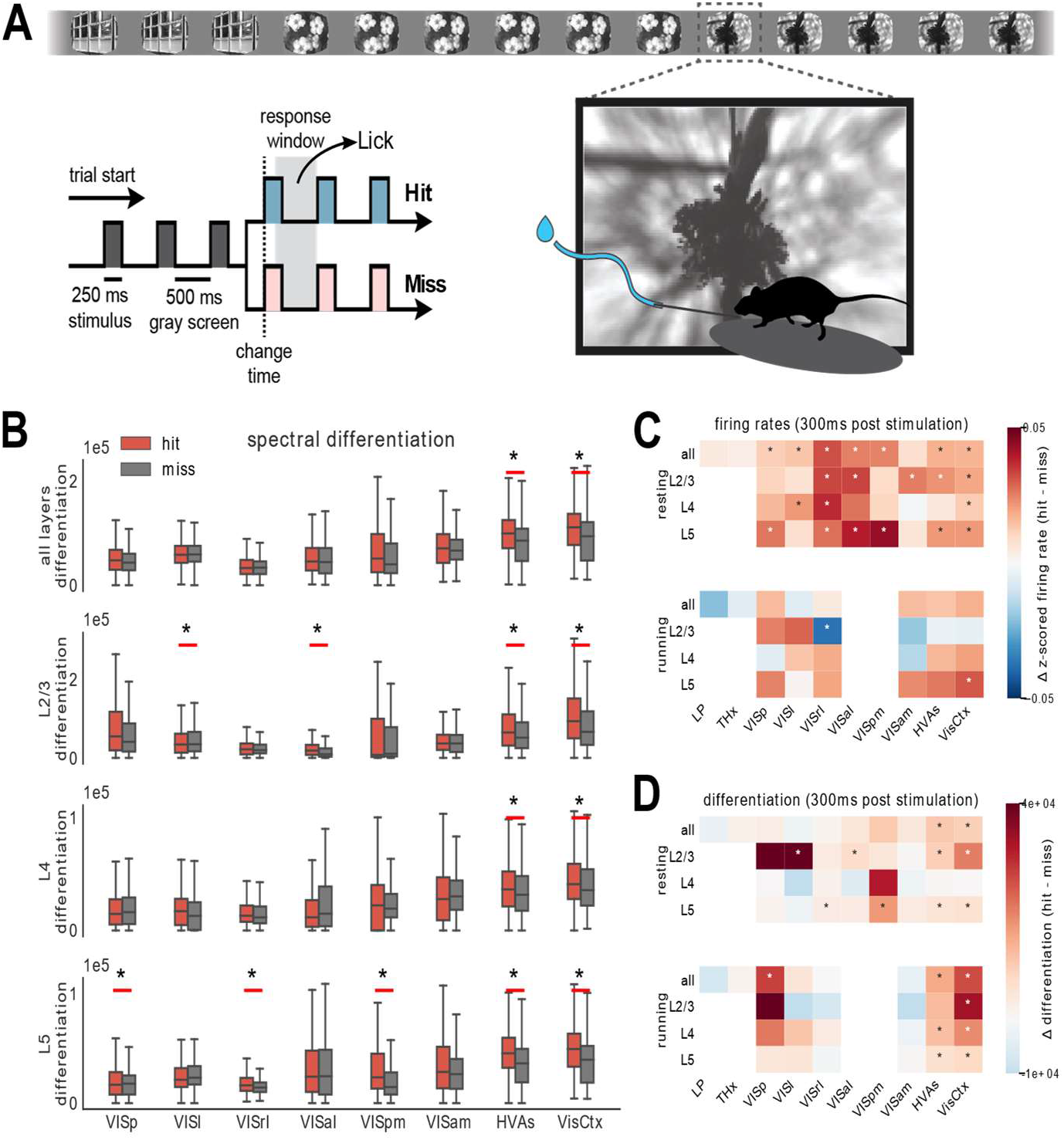
Differentiation during behavior. **A.** Schematic of the change detection task. A series of images is flashed in front of a water-deprived mouse. On detecting a change in the identity of the image, the mouse is trained to lick the spout and is rewarded with water (hit). An undetected change is called a miss. **B.** ND of first 300ms of spiking activity post image presentation for hit (red) and miss trials (grey) across areas and layers. Aggregate areas (all higher visual areas or all visual cortical areas) show a significant difference in ND for hits vs misses, but not individual areas (Benjamini-Hochberg corrected with α = 0.01). Data from single experiment. Summary across all experiments for firing rate **(C)** and ND **(D)** differences between hit and miss. Stars indicate statistically significant differences (Benjamini-Hochberg corrected with α = 0.01). Trials are separated into resting (top) and running (bottom). Differences in ND are stronger when the mouse is running. Hits are associated with elevated firing rate in most areas in the resting state, and almost nowhere in the running state. In contrast, increase in ND is stronger during running, and is specific to aggregate areas.

Indeed, in aggregate ensembles, like all higher visual area neurons (HVAs) or all visual cortical neurons (VisCtx), hits are associated with a more differentiated response in the first 300 ms post stimulus compared to misses (Fig. 5B; significance determined after applying the Benjamini-Hochberg correction with α = 0.01). This difference is statistically significant irrespective of the layer. The 300 ms window is chosen as a trade-off between using the shortest time post-stimulation and including sufficient number of state windows to compute ND while keeping the state lengths close to 100 ms (it also includes the reaction time of the mice). We use state lengths of 60 ms instead of 100 ms for a total of 5 states (300 ms total window length) over which ND is computed. In contrast to the aggregate areas, individual areas do not show a significant difference in ND for hits compared to misses.

In general, hit trials are associated with elevated overall firing rates (Fig. 5D). The increase of firing rates during hits is observed in many individual cortical areas and layers, and in aggregated areas as well. Moreover, the elevation of firing rates is significant largely during trials when the animal is not running, *i.e*., resting, but not as much when the mouse is running (Fig. 5D). Yet, the difference in ND between hits and misses is enhanced when the mouse is running (Fig. 5C). Unlike firing rate, difference in ND for hits and misses appears only in aggregate areas and is not observed in individual areas. ND across regions is thus not a mere reflection of firing activity but appears to be consistent with perception.

### Comparison of ND patterns in the mouse brain visual areas and Convolutional Neural Networks

Like visual areas in the brain, artificial convolutional neural networks (CNNs) also process visual stimuli, although there are fundamental differences in both the structure and function of CNNs and biological neural networks. We apply differentiation analysis to CNNs, treating the activations of units in different CNN layers as neural activity to see how these structural and functional differences are reflected in differentiation.

We analyze three CNN architectures in this study – VGG16 (Simonyan and Zisserman, 2015), Inception V3 (Szegedy et al., 2015) and ResNet50 (He et al., 2015). We use CNN models pre-trained on the ImageNet dataset in Keras (Chollet, 2015). CNNs work on single images and do not have any temporal aspect. However, here we apply the CNN to each frame of the movie, just as the mouse sees it, thus enabling us to extract temporal changes in the activations of different units in the CNN. The CNN classifies the frame into one of the 1000 ImageNet categories. 100 units are subsampled from each layer of the CNN to match the approximate number of neurons recorded from each area in the mouse brain. The timeseries of activations of this fixed subsample of units are used to compute differentiation using the same algorithm as used for the mouse neural activity (see Methods for details).

We see a few key differences in the patterns of differentiation between CNNs and the mouse brain (Fig. 6). Overall, differentiation magnitudes are lower in CNNs (10^4^ - 10^6^) compared to the mouse brain (~10^6^). However, while differentiation changes by a factor of 2 to 3 along the mouse visual hierarchy, it changes by a factor of up to 100 across CNN layers. Unlike the mouse visual cortical areas, for which differentiation remains relatively constant, it increases with depth in CNNs up to an intermediate depth before decreasing again (see Fig. S8 for significance). This combination of differences – lower magnitude, larger range across the hierarchy, and monotonicity up to a certain depth – are seen across all three CNN architectures that we study (Fig. 6B).

**Figure 6:**
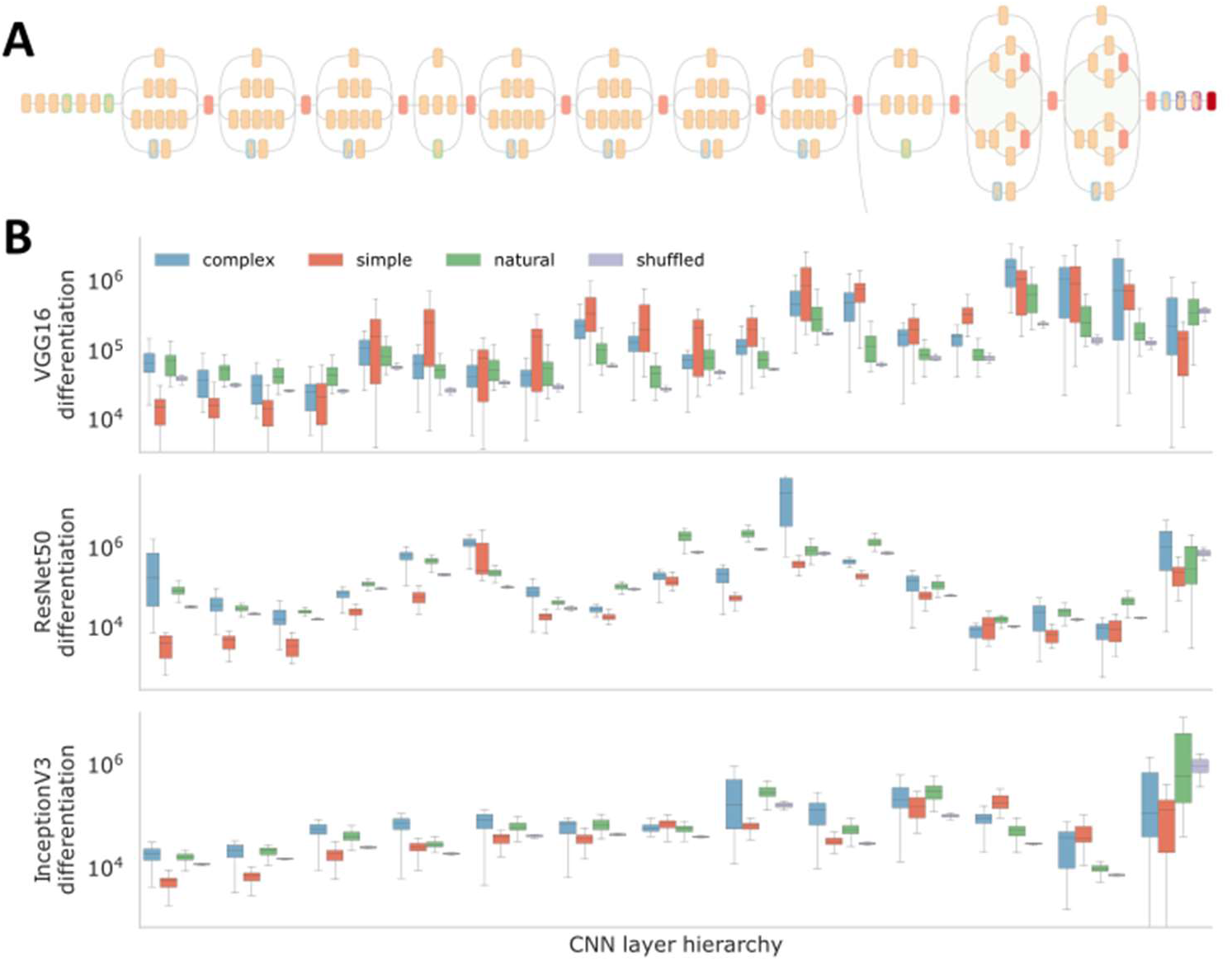
Comparison of differentiation patterns between mouse brain visual areas and Convolutional Neural Networks (CNN). **A.** Schematic of a CNN (Inception v3 pre-trained on the ImageNet database for a 1000-way classification). Units from different layers were subsampled and treated as neurons to compute differentiation in CNNs in response to the same stimuli seen by mice. **B.** Differentiation of activations as a function of depth in the network for the 5 classes of stimuli. Unlike the mouse brain, differentiation increases along the hierarchy, changing by a large factor. Gratings (blue) often evoke higher differentiation compared to natural movies (green), unlike in the mouse brain. This could reflect the fact that the CNN is making a forced classification choice, whereas the mouse visual system is not specialized for that specific task. Other key differences include the fact that the CNN is purely feedforward while the mouse brain is feedback-dominated.

Another key difference is in terms of modulation of differentiation by the stimulus categories, possibly driven by functional differences between the two systems. In the mouse brain, differentiation at the level of entire visual areas (combining all layers) behaves consistently with our expectation based on putative meaningfulness - simple artificial stimuli have the least differentiated response, followed by complex artificial stimuli and finally natural movies. But in CNNs, it is often the case that complex artificial stimuli evoke a higher differentiation than natural movies. The CNNs are trained on a task to classify images into one of 1000 predefined categories. Thus, for example, they often classify slightly different frames of similarly oriented gratings into very different categories. Consequently, a movie of a drifting grating will result in a very highly differentiated response in deeper layers of the CNN, unlike the mouse brain, which likely treats slightly different frames of a grating as equivalent, resulting in lower differentiation compared to a natural movie. Differences in the function of the two systems thus leads to differences in patterns of differentiation with respect to our stimuli.

## Discussion

Differentiation of neural activity at the whole-brain scale in response to subjectively meaningful and meaningless visual stimuli has been studied in humans, although restricted to non-invasive modalities such as fMRI and EEG (Boly et al., 2015; Mensen et al., 2018, 2017) that limit the spatiotemporal resolution of observation of neural activity. This is the first study to characterize the spectral differentiation metric at the cellular level and at the millisecond timescale, for ensembles spanning across multiple visual cortical areas within the mouse brain. Overall, we see that ND of activity in many visual brain regions, and especially at the scale of the entire visual cortex, does not simply reflect stimulus differentiation, but is correlated with the putative meaningfulness of stimuli, consistent with human studies (Fig. 3).

Although, in general, putatively more meaningful stimuli evoke more differentiated neural responses, our analysis revealed a curious exception: in some areas, the no-stimulus condition, corresponding to a grey screen, has a higher ND compared to simple artificial stimuli like flashes and Gabor patches. It is well known that even in the absence of stimuli, structured patterns of activity are spontaneously generated in the brain, often correlated with behavior (Ringach, 2009; Stringer et al., 2019). It is possible that these patterns are more differentiated than the activity evoked by strong but temporally simple external stimuli like full-field flashes, that may strongly constrain activity patterns in the visual system. Temporally more complex inputs (such as gratings or natural movies), on the other hand seem to drive activity across more varied states compared to spontaneous patterns in all the brain regions we investigated. Furthermore, when the mouse is running, simple artificial stimuli evoke a more differentiated response compared to spontaneous activity. This could be because activity in the visual areas is more strongly driven by visual inputs in the running state and spontaneous fluctuations are suppressed (Dadarlat and Stryker, 2017). This is consistent with the interpretation of ND as not just a measure of the passive representation of external information.

Differentiation of activity of entire areas is generally consistent with the putative meaningfulness of stimuli. However, this is not true within individual layers. Though differentiation is significantly modulated by stimuli in all layers, the differentiation patterns in layers 2/3 and 6 are very different from those of other layers or the entire areas; especially with respect to the comparison between complex artificial, natural, and shuffled stimuli. In the behavioral study, we see that differentiation in individual areas is not significantly different for hit and miss trials, but when combined, the aggregate of all higher visual areas shows significantly higher ND for hits compared to misses (Fig. 5). Together these observations suggest that the activity in the visual cortex as a whole might be more correlated with meaningfulness than in its individual parts.

In the canonical cortical microcircuit, feedforward information propagates upwards from L2/3 (Bastos et al., 2012), while L5 pyramidal (L5p) cells are involved in integrating feedback and feedforward streams of information, mediated by thalamic inputs (Aru et al., 2020; Suzuki and Larkum, 2020), and propagating outputs to other cortical and subcortical areas. Recent work suggests that such dendritic integration of information by L5p cells is required for experiencing specific contents of consciousness (Takahashi et al., 2016). Consistent with this picture, in our study also, we observe that differentiation is correlated with putative meaningfulness in deep layers that integrate feedback and feedforward information, mediated by the thalamus; but not in layer 2/3, which may be related to the larger role of this layer in relaying feedforward information to higher areas.

In a passive viewing paradigm, it is not possible to ascertain the ground truth regarding the perception of animals. To address this, we analyzed ND in a behavioral paradigm, where there is higher certainty regarding the perception of animals (Fig. 5). The mice might be following different strategies in the change detection task, some of which not involving perception of the images: for example, they could learn the average time between image changes and lick randomly around those times. For such strategies, we do not expect any difference in ND for hits or misses. Yet, we find that hits correspond to significantly higher ND in aggregate areas (HVAs or the entire visual cortex) compared to misses, consistent with the intended strategy - perceived images changes lead to hits and unperceived ones to misses. Moreover, although hits evoked higher firing rates compared to misses in the resting state in many individual areas, differences in ND were restricted to aggregate areas, reflecting the specificity of the ND metric.

A comparison between differentiation in the mouse brain and CNNs reveals interesting differences (Fig. 6). Overall, the dynamic range of differentiation across the hierarchy, even after normalizing for the mean firing rate (or activations), is much higher for CNNs than the mouse brain. Secondly, differentiation increases with depth up to an intermediate layer in the CNN hierarchy. Recent insights into the mechanism of CNNs trained for image classification show that these networks initially expand the dimensionality of representations by generating features, and the final layers select low-dimensional combinations of these features to ultimately classify inputs into a few output categories (Ansuini et al., 2019; Cohen et al., 2020; Recanatesi et al., 2019). Differentiation trends thus reflect the trends in dimensionality of representations, peaking in intermediate layers. This is not unexpected given that the distances between state vectors should increase with increasing embedding dimensionality. In the mouse visual system also, the dimensionality of evoked neural activity increases along the hierarchy, indicating a similar expansion of feature representations (Dahmen et al., 2020). However, ND of neural activity, purportedly reflecting perception and not just representation, remains constant over the mouse visual hierarchy. Some other differences between CNNs and mouse brains might be arising out of functional differences between the two systems – as discussed earlier, since the CNN is sensitive to small changes in the input images and is performing a forced choice classification task, stimuli such as gratings evoke higher differentiation than natural movies in CNNs unlike the mouse brain.

The notion of differentiation is grounded in the 1-to-1 mapping between subjective perception and its neural representation. Temporal differentiation of activity is also postulated by the integrated information theory of consciousness (IIT) as a necessity for conscious experience, in addition to integration (activity in different parts or at different times should not be completely independent) (Koch et al., 2016; Ruiz de Miras et al., 2019; Sarasso et al., 2014; Tononi et al., 2016). According to this theory, a combined measure of integration and differentiation, Phi, is expected to be maximal at specific spatial and temporal scales, corresponding to the spatiotemporal scale of experience (Tononi et al., 2016). As we observe, just the differentiation (without integration) of ensemble activity does not have an optimal timescale, but we do find an optimal timescale of ~100 ms for single neuron differentiation (Fig. 2).

In conclusion, we find that Neurophysiological Differentiation can be a useful tool to identify specific subpopulations of neurons that may be involved in subjective perception. These results, in conjunction with human studies, where ND is found to correlate with subjective reports of meaningfulness, reflect the future potential to objectively infer the quantity of experience of subjects who otherwise have no ability to report it, such as animals or humans with disorders of consciousness.

## Methods

### Computing spectral differentiation

Spectral differentiation is computed on timeseries data. Therefore, for each neuron, spike times are converted into a binary timeseries with a resolution of 5 ms bins (200 Hz), depending on the presence or absence of a spike in each bin. This binary timeseries is then convolved with a normalized Gaussian kernel with a halfwidth of 2 bins (10 ms) and cut off on either side at 5 bins (25 ms) to obtain the smoothened firing rates.

Spectral Differentiation is defined for an ensemble of *N* neurons over windows of size *W*, for states with state length *S* (*W* / *S* state intervals in total per window). For each state interval, the ‘spectral state’ of that interval is quantified in terms of the power spectrum of the signal of each neuron within the state window: for this, the spectra are first computed using *numpy.fft.rfftn* and the absolute value is squared to obtain the power spectrum. Since the sampling frequency is fixed at 200 Hz, power spectrum is obtained over frequencies ranging from −100 to 100 Hz with a resolution of 1/*S* Hz. The power spectra for each neuron are concatenated into a (200 *N* / *S*) dimensional vector. Euclidean distance is computed between all pairs of such *W*/*S* vectors and the median distance is defined as the spectral differentiation.

### Normalization of spectral differentiation

Spectral differentiation is an unbounded measure, and the absolute value of the measure is not directly meaningful. The relative magnitude of the metric across different stimuli, or across different ensembles is of primary interest.

To enable comparison of the metric across different ensembles of neurons, a few normalizations are performed. First, simply scaling all firing rates by a constant factor Q changes ND by a factor of Q^2^. To account for the differences in the mean firing rates, the firing rates are divided by the ensemble mean firing rate before computing differentiation.

Second, as the number of neurons increases, the dimensionality of the state space in which distances are computed also increases. To account for the variations in the number of neurons across experiments and brain regions, we normalize by an additional factor of *sqrt(N)*, where *N* is the number of neurons.

The state length also affects the dimensionality of the final state vector, which is given by 200 *N/S*. We correct for this by multiplying differentiation by *sqrt(S)*. Furthermore, the total power in FFT scales as the square of the number of samples, i.e., state length, so differentiation is further divided by *S^2^*.

Overall, we have

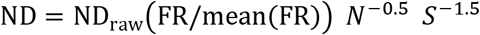

Where ND_raw_(·) is as described in the preceding section (computing spectral differentiation) and FR is the vector of time-binned firing rates for all neurons in the analyzed ensemble.

### Characteristic timescale of stimuli and neural activity

‘Characteristic timescale’ is the timescale over which the signal changes substantially. This is simply the time for which the signal remains constant before changing for piecewise constant uncorrelated signals such as the toy example of Fig. 2A (600 ms), or stimuli like Gabors, static gratings (250 ms) and shuffled movies (33 ms), which change after a fixed period of time and take random values. The Flashes stimulus remains grey for 1.75 s and turns black or white for 0.25 s, and thus has a characteristic timescale of 1 s on average over which it remains constant.

For all other signals, the characteristic timescale is the timescale over which the autocorrelation of the signal decays exponentially. To obtain autocorrelation timescales (AC), autocorrelation of firing rates (or pixel intensities for stimulus AC) are first computed. The autocorrelation is best modeled by two exponents, over an initial fast timescale and a slower timescale, with the transition point between the timescales changing from neuron to neuron. To account for this, the logarithm of the autocorrelation is fit using *scipy.curvefit* to:

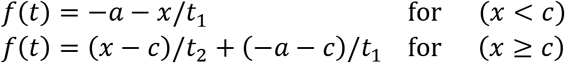

Parameters *a, c, t_1_, t_2_* are fit to data. *t_1_* is constrained between 0 and 1.5 s; *t_2_* is constrained between 0.5 and 20 s; and *c* between 0.03 s and 0.2 s. We only consider the fast timescale *t_1_* for all other analysis.

Note that for piecewise constant signals described earlier, the autocorrelation does not give a good estimate of the timescale over which the signal typically remains constant.

### Convolutional Neural Networks

Pre-trained models of three CNNs (VGG16, ResNet50 and InceptionV3) from the Python package Keras are used for this analysis. The networks are pre-trained on an image classification task using the ImageNet database.

For stimuli such as flashes, Gabors, static and drifting gratings and the shuffled movie, sample movies of length 90 s are generated following the same parameters as those presented to the mice. Natural movies are used as is. All movies are sampled at 200 FPS.

For a given CNN model, 100 units from each layer are selected randomly and fixed for subsequent analysis, akin to sampling a few fixed neurons across the mouse visual cortex. The CNN model is run on each frame of a movie, and the corresponding activations from the pre-selected units from each layer are recorded. We thus obtain a 90 s long timeseries sampled at 200 Hz (same as the sampling time of firing rates) for each unit. These are then used to compute ND of activity in each CNN layer separately as a function of time (following **Methods: computing spectral differentiation**). In this way, although CNNs do not have an inherent time dependence, we can compute the temporal complexity of CNN responses to time-varying stimuli. Stimuli are then grouped into simple artificial, complex artificial, natural, and shuffled, and ND samples from each group are plotted as a box plot for Fig. 6.

## Acknowledgements

We thank the Allen Institute founder, Paul G. Allen, for his vision, encouragement, and support; Allan Jones for providing the crucial environment that enabled our large-scale team effort; Stefan Mihalas, Leslie Claar, Ethan McBride, Irene Rembado and Dana Mastrovito for helpful discussions.

## References

1. Ansuini A, Laio A, Macke JH, Zoccolan D (2019) Intrinsic dimension of data representations in deep neural networks. ArXiv190512784 Cs Stat.

2. Aru J, Suzuki M, Larkum ME (2020) Cellular Mechanisms of Conscious Processing. Trends Cogn Sci 24:814–825.

3. Barttfeld P, Uhrig L, Sitt JD, Sigman M, Jarraya B, Dehaene S (2015) Signature of consciousness in the dynamics of resting-state brain activity. Proc Natl Acad Sci 112:887–892.

4. Bastos AM, Usrey WM, Adams RA, Mangun GR, Fries P, Friston KJ (2012) Canonical microcircuits for predictive coding. Neuron 76:695–711.

5. Boly M, Sasai S, Gosseries O, Oizumi M, Casali A, Massimini M, Tononi G (2015) Stimulus Set Meaningfulness and Neurophysiological Differentiation: A Functional Magnetic Resonance Imaging Study. PLOS ONE 10:e0125337.

6. Brette R (2019) Is coding a relevant metaphor for the brain? Behav Brain Sci 42.

7. Buzsáki G (2019) The brain from inside out. New York, NY: Oxford University Press.

8. Chollet F (2015) Keras. Keras.

9. Cohen U, Chung S, Lee DD, Sompolinsky H (2020) Separability and geometry of object manifolds in deep neural networks. Nat Commun 11:746.

10. Dadarlat MC, Stryker MP (2017) Locomotion Enhances Neural Encoding of Visual Stimuli in Mouse V1. J Neurosci 37:3764–3775.

11. Dahmen D, Recanatesi S, Jia X, Ocker GK, Campagnola L, Jarsky T, Seeman S, Helias M, Shea-Brown E (2020) Strong and localized coupling controls dimensionality of neural activity across brain areas (preprint). Neuroscience.

12. Del Cul A, Baillet S, Dehaene S (2007) Brain dynamics underlying the nonlinear threshold for access to consciousness. PLoS Biol 5:e260.

13. Gosseries O, Schnakers C, Ledoux D, Vanhaudenhuyse A, Bruno M-A, Demertzi A, Noirhomme Q, Lehembre R, Damas P, Goldman S, Peeters E, Moonen G, Laureys S (2011) Automated EEG entropy measurements in coma, vegetative state/unresponsive wakefulness syndrome and minimally conscious state. Funct Neurol 26:25.

14. He K, Zhang X, Ren S, Sun J (2015) Deep Residual Learning for Image Recognition. ArXiv151203385 Cs.

15. Herzog MH, Drissi-Daoudi L, Doerig A (2020) All in Good Time: Long-Lasting Postdictive Effects Reveal Discrete Perception. Trends Cogn Sci 24:826–837.

16. Herzog MH, Kammer T, Scharnowski F (2016) Time Slices: What Is the Duration of a Percept? PLoS Biol 14:e1002433.

17. Hudetz AG, Liu X, Pillay S (2015) Dynamic Repertoire of Intrinsic Brain States Is Reduced in Propofol-Induced Unconsciousness. Brain Connect 5:10–22.

18. Koch C (2004) The Quest for Consciousness.

19. Koch C, Massimini M, Boly M, Tononi G (2016) Neural correlates of consciousness: progress and problems. Nat Rev Neurosci 17:307–321.

20. Mayner WGP et al. (2022) Measuring stimulus-evoked neurophysiological differentiation in distinct populations of neurons in mouse visual cortex. eNeuro. doi 10.1523/ENEURO.0280-21.2021

21. Mensen A, Marshall W, Sasai S, Tononi G (2018) Differentiation Analysis of Continuous Electroencephalographic Activity Triggered by Video Clip Contents. J Cogn Neurosci 30:1108–1118.

22. Mensen A, Marshall W, Tononi G (2017) EEG Differentiation Analysis and Stimulus Set Meaningfulness. Front Psychol 8:1748.

23. Piasini E, Soltuzu L, Muratore P, Caramellino R, Vinken K, Op de Beeck H, Balasubramanian V, Zoccolan D (2021) Temporal stability of stimulus representation increases along rodent visual cortical hierarchies. Nat Commun 12:4448.

24. Recanatesi S, Farrell M, Advani M, Moore T, Lajoie G, Shea-Brown E (2019) Dimensionality compression and expansion in Deep Neural Networks. ArXiv190600443 Cs Stat.

25. Ringach DL (2009) Spontaneous and driven cortical activity: implications for computation. Curr Opin Neurobiol 19:439–444.

26. Rolls ET (2000) Functions of the Primate Temporal Lobe Cortical Visual Areas in Invariant Visual Object and Face Recognition. Neuron 27:205–218.

27. Ruiz de Miras J, Soler F, Iglesias-Parro S, Ibáñez-Molina AJ, Casali AG, Laureys S, Massimini M, Esteban FJ, Navas J, Langa JA (2019) Fractal dimension analysis of states of consciousness and unconsciousness using transcranial magnetic stimulation. Comput Methods Programs Biomed 175:129–137.

28. Sarasso S, Rosanova M, Casali AG, Casarotto S, Fecchio M, Boly M, Gosseries O, Tononi G, Laureys S, Massimini M (2014) Quantifying Cortical EEG Responses to TMS in (Un)consciousness. Clin EEG Neurosci 45:40–49.

29. Shimojo S, Paradiso M, Fujita I (2001) What visual perception tells us about mind and brain. Proc Natl Acad Sci 98:12340–12341.

30. Siegle JH et al. (2021) Survey of spiking in the mouse visual system reveals functional hierarchy. Nature 592:86–92.

31. Simonyan K, Zisserman A (2015) Very Deep Convolutional Networks for Large-Scale Image Recognition. ArXiv14091556 Cs.

32. Solovey G, Alonso LM, Yanagawa T, Fujii N, Magnasco MO, Cecchi GA, Proekt A (2015) Loss of Consciousness Is Associated with Stabilization of Cortical Activity. J Neurosci 35:10866–10877.

33. Stringer C, Pachitariu M, Steinmetz N, Reddy CB, Carandini M, Harris KD (2019) Spontaneous behaviors drive multidimensional, brainwide activity. Science 364.

34. Suzuki M, Larkum ME (2020) General Anesthesia Decouples Cortical Pyramidal Neurons. Cell 180:666–676.e13.

35. Synek VM (1988) Prognostically important EEG coma patterns in diffuse anoxic and traumatic encephalopathies in adults. J Clin Neurophysiol Off Publ Am Electroencephalogr Soc 5:161–174.

36. Szegedy C, Vanhoucke V, Ioffe S, Shlens J, Wojna Z (2015) Rethinking the Inception Architecture for Computer Vision. ArXiv151200567 Cs.

37. Takahashi N, Oertner TG, Hegemann P, Larkum ME (2016) Active cortical dendrites modulate perception. Science 354:1587–1590.

38. Tononi G, Boly M, Massimini M, Koch C (2016) Integrated information theory: from consciousness to its physical substrate. Nat Rev Neurosci 17:450–461.

