## Supplemental figures and tables for "A survey of neurophysiological differentiation across mouse visual brain areas and timescales"

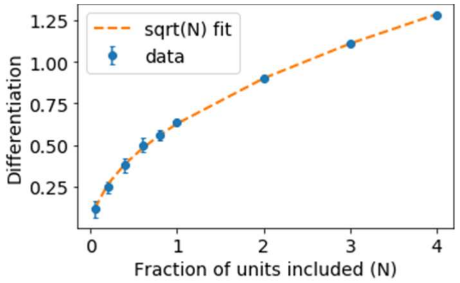


**Figure S1: Normalizations applied to differentiation. *A***. *Differentiation scales with the square root of number of neurons. Different fractions of neurons were subsampled from a large population of neurons from a single experiment, and spectral differentiation was computed for each sample (blue points with SD shown as error bars). Orange line shows a sqrt(N) fit.*

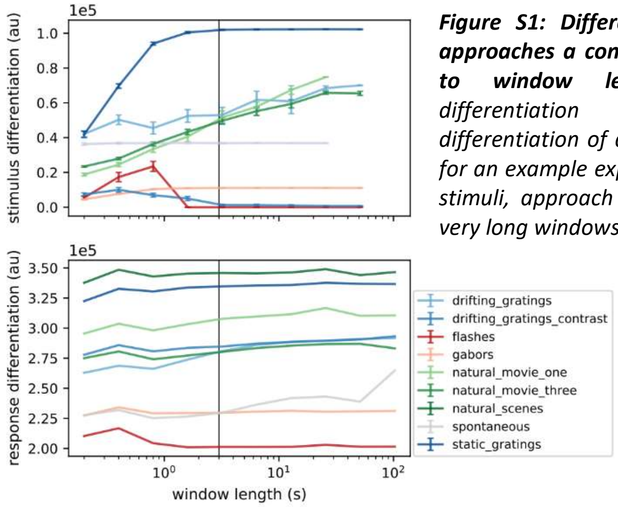


***Figure S1: Differentiation asymptotically approaches a constant value with respect to window length.** Both stimulus differentiation (top) and response differentiation of all visual cortical neurons for an example experiment (bottom), for all stimuli, approach an asymptotic value for very long windows.*

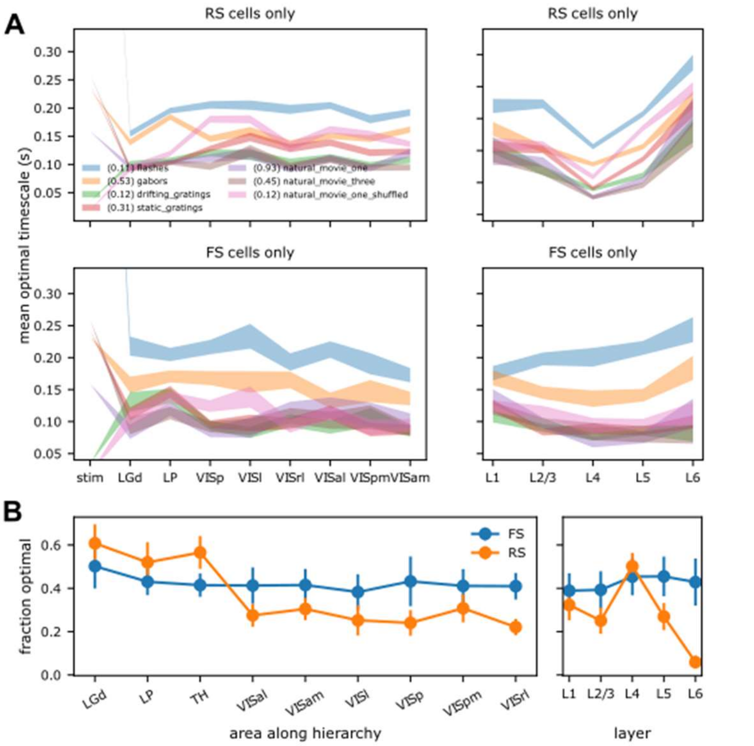


***Figure S3: Optimal timescale for neurons – dependence on stimuli and brain region. A.** Mean and standard deviation (shaded region) of the optimal timescale for individual regular spiking (RS) and fast spiking (FS) neurons in Neuropixels recordings from the different brain areas or cortical layers. Areas are arranged along the x-axes according to the level in the anatomical hierarchy (Harris et al., Nature, 2019). The optimal timescale increases from thalamus to the cortex, but remains constant throughout the cortex for RS cells; and decreases going up the hierarchy for FS cells. Across cortical layers, optimal timescale of RS neurons decreases with depth, with the minimum at L5; and does not vary much with layer for FS neurons. Optimal timescales for neurons are unrelated to those for stimuli (leftmost points). **B.** Fraction of neurons in each brain region that have a physiologically relevant optimal timescale (>10 ms). The fraction of FS or RS neurons with an optimal timescale is highest in the thalamic areas, and much reduced in cortical or hippocampal areas. Within the cortex, this fraction is highest in layers 4 and 5 for RS neurons. Error bars indicate SD across different stimuli*

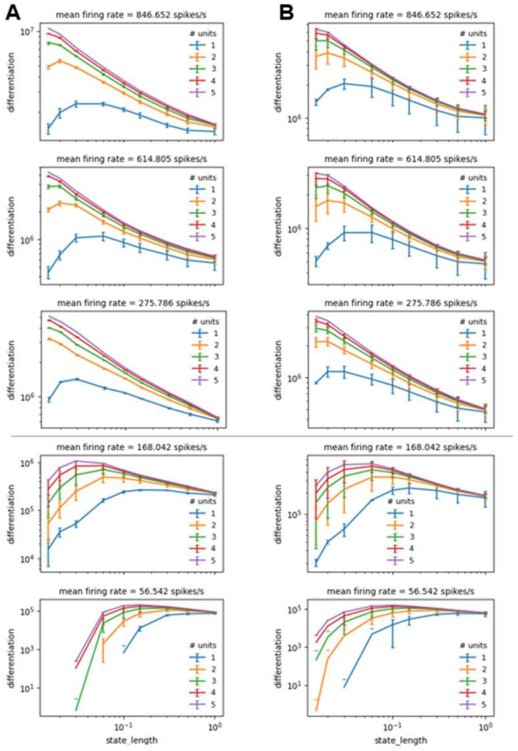


***Figure S4**: **Jitter or shift in firing times of high firing neurons causes loss of optimal timescale in ensembles.** Constant temporal shift or jitter was added to the spike times of 5 neurons with different average firing rates. For each neuron, 5 copies of the spike trains were generated with either a shift of between -10 and 10 ms (**A**) or jitter of up to 5 ms (**B**) introduced randomly in each case. Each of the 5 altered timeseries were treated as distinct pseudoneurons, and differentiation as a function of timescale is plotted for combinations of 1 through 5 pseudoneurons. For neurons with high firing rates (top 3 panels), the optimal timescale becomes physiologically irrelevant (<10 ms) at the ensemble level. For low firing neurons (bottom 2 panels), optimality of timescales is retained although shifted to a shorter time.*

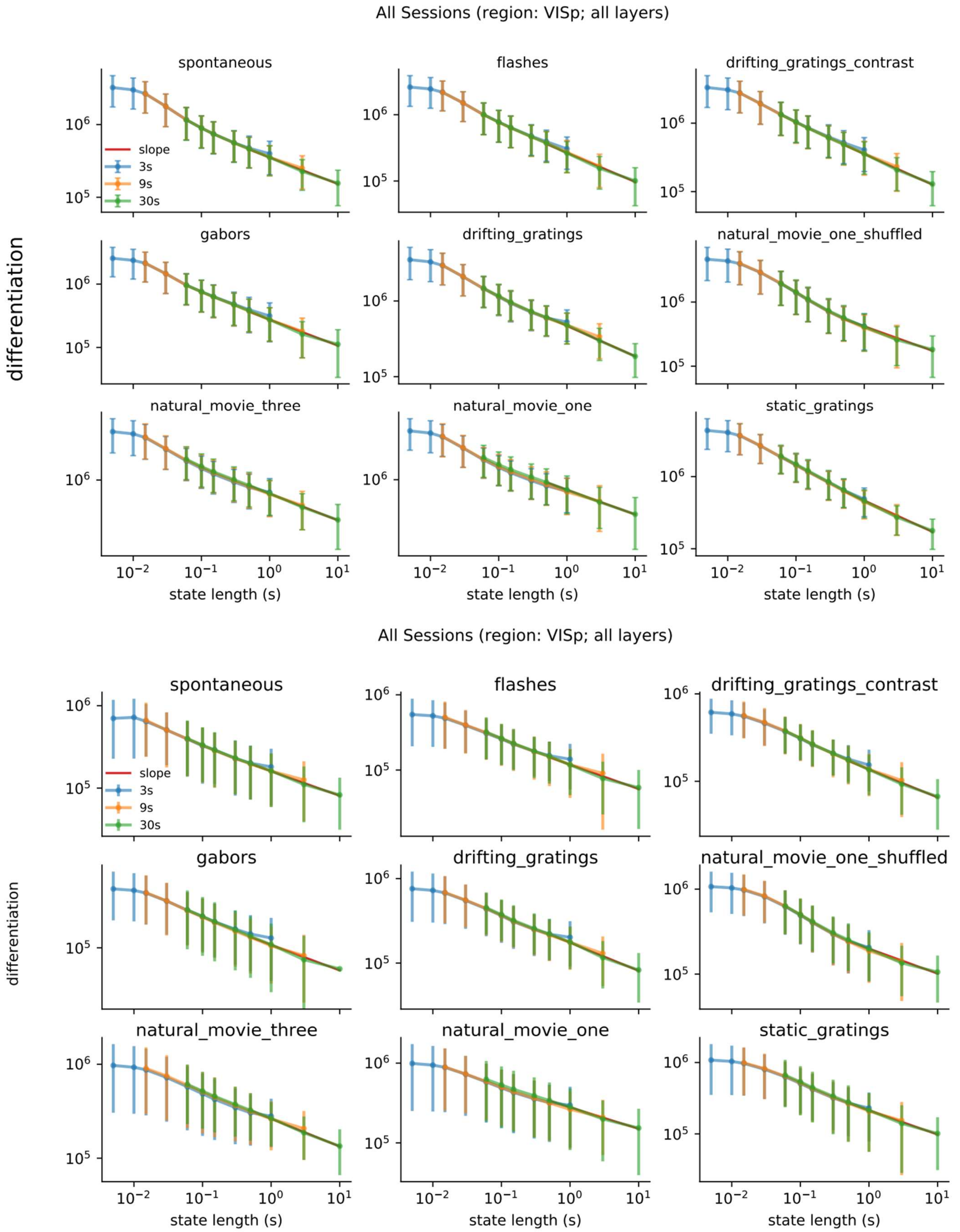


**Figure S5: ND does not have an optimal timescale between 10 ms to 10 s for all stimuli. Top.** Regular spiking units. **Bottom.** Fast spiking units.

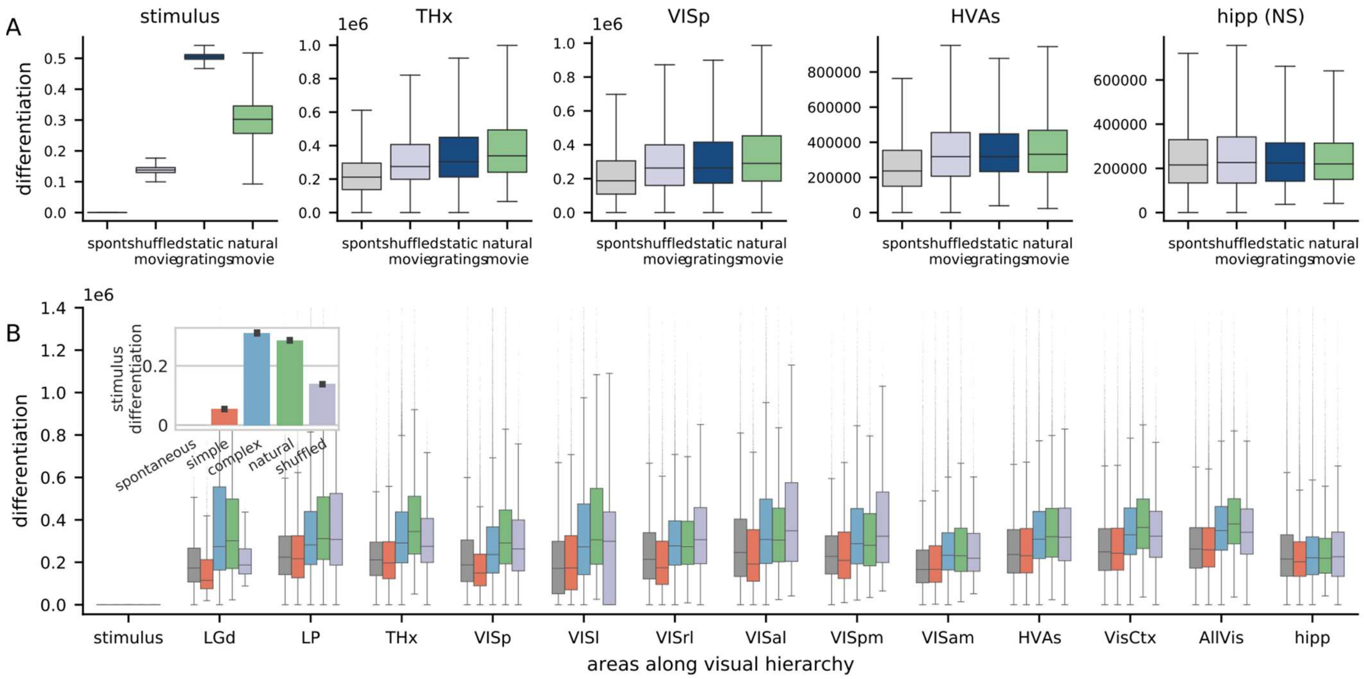


**Figure S6: Area-wise modulation of differentiation by stimuli for fast spiking neurons. A.** Similar to regular spiking neurons, differentiation of fast spiking neurons is also statistically significantly different ($p < 0.001$) between all pairs of stimuli shown in Thalamus, VISp and higher visual areas. All pairwise differences are however insignificant in the hippocampus. **B.** Note that ND for fast spiking neurons is lower than regular spiking neurons by roughly a factor of 2, but the modulation by stimuli is largely unchanged. Secondly, magnitude of ND in thalamic regions is unchanged between FS and RS neurons (see Fig. 3 for comparison).

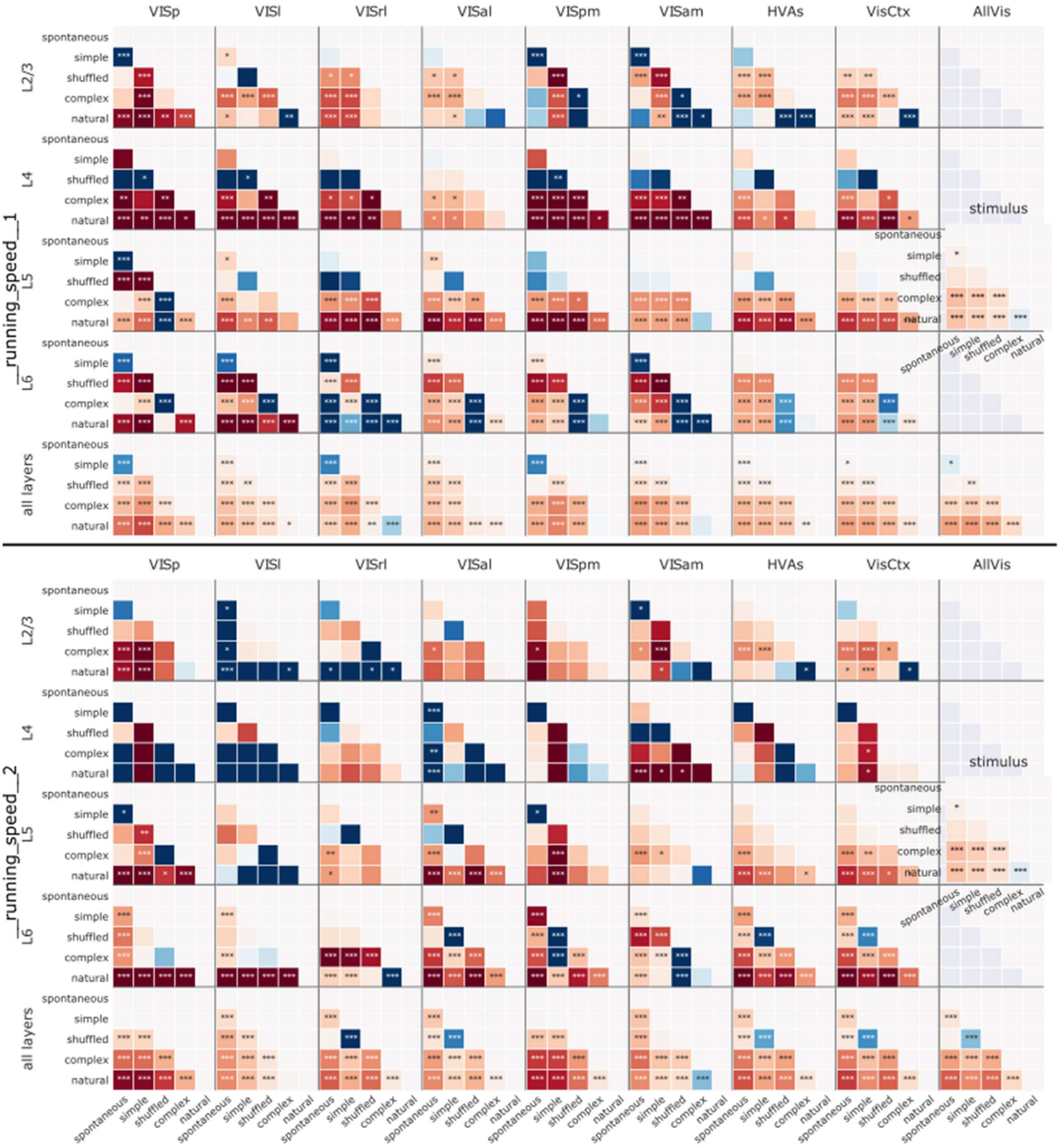


**Figure S7: Modulation of differentiation by stimuli across different areas and layers in running and resting states.** Running speed 1 (top) is the resting state and running speed 2 (bottom) is running state of the animal. While resting, shuffled movies evoke higher differentiation compared to natural movies in layer 6; but not while the mouse is running. Differentiation modulation by stimuli is much less significant in layers 2/3, 4 and 5 when the mouse is running compared to the resting state.

| | stimulus | correlation | p-value |
|---|---|---|---|
| VGG16 (up to 16) | complex | 0.447336 | 2.043038e-48 |
| | natural | 0.422617 | 5.353827e-36 |
| | shuffled | 0.604647 | 2.527264e-17 |
| | simple | 0.252380 | 2.319659e-22 |
| ResNet50 (up to 11) | complex | 0.082606 | 3.385512e-02 |
| | natural | 0.671780 | 1.925059e-73 |
| | shuffled | 0.725309 | 3.225824e-19 |
| | simple | 0.122419 | 1.127009e-04 |
| InceptionV3 (up to 10) | complex | 0.486563 | 5.526303e-37 |
| | natural | 0.486991 | 3.878880e-31 |
| | shuffled | 0.601209 | 3.743691e-11 |
| | simple | 0.517061 | 1.117878e-62 |

| | stimulus | correlation | p-value |
|---|---|---|---|
| VGG16 (16 onwards) | complex | 0.323536 | 2.982855e-07 |
| | natural | -0.011031 | 8.767979e-01 |
| | shuffled | 0.107171 | 5.104004e-01 |
| | simple | 0.021042 | 6.907103e-01 |
| ResNet50 (11 onwards) | complex | -0.465983 | 8.346108e-21 |
| | natural | -0.677327 | 1.275326e-41 |
| | shuffled | -0.839657 | 5.189094e-17 |
| | simple | -0.596337 | 2.623994e-53 |
| InceptionV3 (10 onwards) | complex | -0.674080 | 3.369051e-25 |
| | natural | -0.752189 | 1.341123e-28 |
| | shuffled | -0.944722 | 4.336741e-15 |
| | simple | -0.333435 | 1.965870e-08 |

***Fig. S8: Significant increase of differentiation up to an intermediate layer in CNNs followed by a decrease***
